# *Mettl14* is required for mouse post-implantation development by facilitating epiblast maturation

**DOI:** 10.1101/294926

**Authors:** Tie-Gang Meng, Xukun Lu, Lei Guo, Guan-Mei Hou, Xue-Shan Ma, Qian-Nan Li, Lin Huang, Li-Hua Fan, Zheng-Hui Zhao, Xiang-Hong Ou, Ying-Chun OuYang, Heide Schatten, Lei Li, Zhen-Bo Wang, Qing-Yuan Sun

## Abstract

N6-methyladenosine (m^6^A) is the most prevalent and reversible internal modification of mammalian messenger and noncoding RNAs mediated by specific m^6^A writer, reader, and eraser proteins. As an m^6^A writer, the METTL3–METTL14-WTAP complex dynamically regulates m^6^A modification and plays important roles in diverse biological processes. However, our knowledge about the complete functions of this RNA methyltransferase complex, the contributions of each component to the methylation and their impacts on different biological pathways, are still very limited. Here, by employing both *in vivo* and *in vitro* models, we report that METTL14 was indispensable for post-implantation embryonic development by facilitating the conversion from naïve to primed state of the epiblast. Depletion of *Mettl14* lead to conspicuous embryonic growth retardation from E6.5 mainly as a result of resistance to differentiation, which further lead to embryonic lethality early in gestation. Our data highlight the critical function of METTL14, as an m^6^A modification regulator, in orchestrating early mouse embryogenesis.

## INTRODUCTION

Akin to the roles of DNA methylation and histone modifications in epigenetics, N6-methyl-adenosine (m^6^A) is the most abundant internal modification of mRNAs in eukaryotes, and dysregulation of this modification has already been clearly linked to many human diseases, such as obesity, cancer and intellectual disability (Sibbritt et al., 2013). Importantly, since fat-mass and obesity-associated protein (FTO) and ALKBH5 have been shown to act as m^6^A demethylases, it is commonly accepted that m^6^A modifications are reversible in mammalian cells (Jia et al., 2011; Zheng et al., 2013), evoking the biological implications of the dynamic m^6^A modification, which is incompletely understood in mammals.

m^6^A formation is catalyzed by the RNA methyltransferase complex containing methyltransferase like 3 (METTL3), methyltransferase like 14 (METTL14) and Wilms’ tumor 1-associating protein (WTAP), the writer of the m^6^A marks(Wang et al., 2014). It is interesting and fascinating that the RNA methyltransferase complex contains two methyltransferase subunits, which could catalyze m^6^A formation, respectively. Furthermore, it appears that METTL3 and METTL14 impact different targets, in spite of a common set of substrates (Liu et al., 2014). Contributions from each component to the methylation and their impacts on different biological pathways are unclear. METTL3 is the first reported mammalian m^6^A methyltransferase. *Mettl3*-null mutant embryos died by embryonic day 6.5 (E6.5) as a result of failure to down-regulate *Nanog* mRNA level (Geula et al., 2015). However, the function of METTL14 in mammalian development is largely unclear.

Early embryonic development in mammals involves a coordinated cell lineage specification from the pluripotent epiblast to diverse types of cells around implantation, which lays the foundation for a successful body plan (Arnold and Robertson, 2009). During this process, naïve pluripotency markers (*Nanog, Esrrb, Nr5a2, Klf2, Klf4, Rex1*) are down-regulated, and primed pluripotency markers (*Fgf5, Otx2, Sox3, Dnmt3b, Wnt3*) are up-regulated, enabling the conversion of the pluripotent epiblast from the naïve to primed state (Kalkan et al., 2017), a process commonly known as epiblast maturation. This conversion is developmentally essential, failure of which would result in early embryonic lethality (Geula et al., 2015; Huang et al., 2017). *In vitro* models, including embryonic stem cells (ESCs) (Evans and Kaufman, 1981; Martin, 1981), epiblast-like cells (EpiLCs) (Hayashi et al., 2011) and epiblast stem cells (EpiSCs) (Brons et al., 2007; Tesar et al., 2007) that represent different states of pluripotency *in vivo* have provided many insights into the molecular regulation of the process. However, due to the limitation of materials, techniques and inaccessibility to the process *in vivo*, what we know about epiblast maturation and mammalian early embryogenesis is still very limited.

The aim of our present study was to determine the function of METTL14 during embryo development. We deleted *Mettl14* gene using the CRISPR/cas9 system. We found that depletion of *Mettl14* render mouse naïve epiblast or embryonic stem cells resistant to differentiation, leading to embryonic lethality by E6.5.

## RESULTS

### Establishment of a *Mettl14* knockout mouse model

To investigate the physiological function of *Mettl14*, we established a line of *Mettl14* knockout mice using the CRISPR/Cas9 system. Oligos encoding sgRNA that targets the exon2 of *Mettl14* were inserted into px330 plasmid. Unique sgRNA sequences were chosen based on the Genetic Perturbation Platform from the Broad Institute website (Cong et al., 2013). We constructed a *Mettl14* targeting vector and microinjected it with Cas9 mRNA and gRNA into zygotes of C57BL/6 mice. The sgRNA was designed to target the exon 2 of the endogenous mouse *Mettl14* gene. Six *Mettl14^+/-^* mice were obtained from these experiments. The genotype of mice including *Mettl14^+/+^* and *Mettl14^+/-^* were confirmed by DNA sequencing and nucleic acid electrophoresis (Fig. 1).

**Figure 1.**
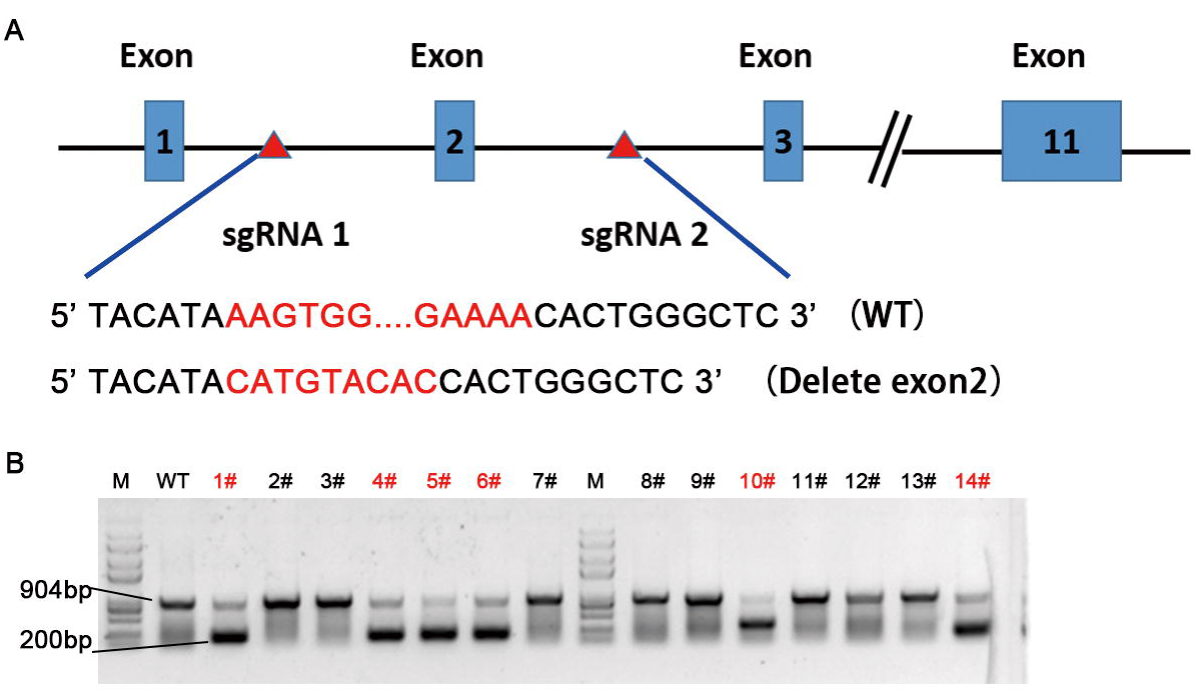
Establishment of *Mettl14* knockout mouse model. (A) Scheme of targeting *Mettl14* using the CRISPR/Cas9 system. Two sgRNAs were designed to target the intron 1 and intron 2, respectively, to delete the exon 2 in *Metttl14*. DNA sequence of the wild type and the resulting mutant alleles were shown. (B) Genotyping result of the founder mice. Six *Mettl14^+/-^* founder mice were obtained.

Thus, we successfully established a *Mettl14* knockout mouse model, which could be used for further phenotypic and functional analysis.

### METTL14 is required for mouse early development

Genotyping of postnatal offspring by PCR analysis from *Mettl14* heterozygous intercrosses revealed the absence of mouse homozygous for the *Mettl14* mutation, indicating possible embryonic lethality of *Mettl14* knockout mice early during gestation. In order to determine when the *Mettl14* mutation produced a lethal phenotype, embryos of heterozygous intercrosses were obtained from E3.5-E12.5 (Fig. 2A). Genotyping of these embryos by PCR analysis revealed that *Mettl14* knockout embryos were observed at Mendelian ratios until E6.5, but no mutants were identified at E8.5, E10.5 and E12.5. Although *Mettl14* knockout embryos could be detected at E6.5 and E7.5, they exhibited largely growth retardation and aberrant morphology (Fig. 2B-2E). Thus, these data suggest that *Mettl14* is indispensable for mouse early development.

**Figure 2.**
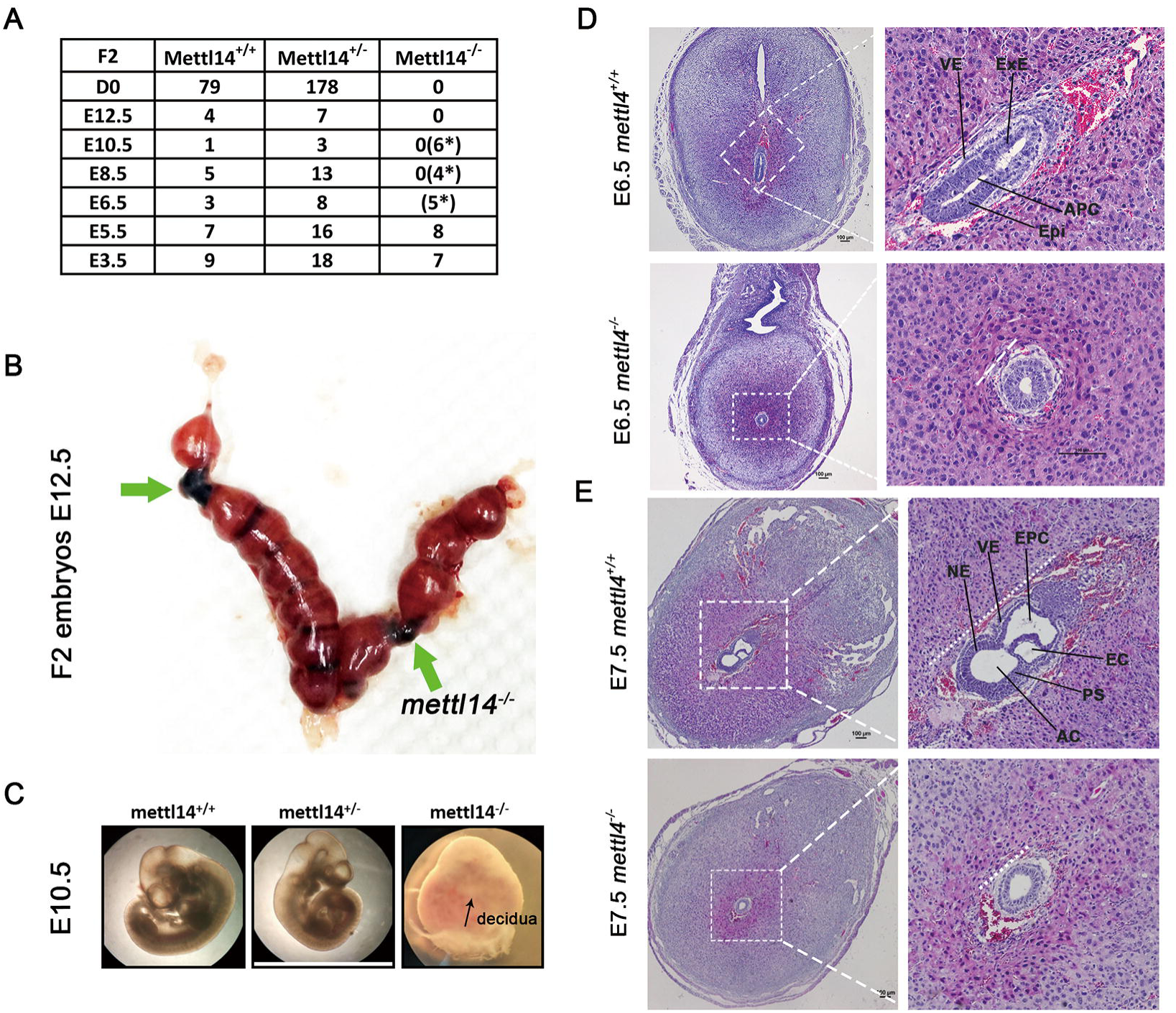
Depletion of *Mettl14* leads to early embryonic lethality. (A) Development of *Mettl14^-/-^* mice. Embryos from *Mettl14^+/-^* intercrosses were obtained at multiple stages, and the genotypes were determined using PCR. The numbers of *Mettl14^+/+^*, *Mettl14^+/-^* and *Mettl14^-/-^* embryos at each stage were shown. (B and C) *Mettl14^-/-^* embryos at E12.5 (B, green arrows) and E10.5 (C) were completely resorbed. (D and E) Histological analysis of control and *Mettl14^-/-^* embryos at E6.5 and E7.5. Note the growth retardation and malformation of E6.5 and E7.5 *Mettl14^-/-^* embryos. VE, visceral endoderm; Epi, epiblast; ExE, extra-embryonic ectoderm; PAC, proamniotic cavity; NE, neuroectoderm; AC, amniotic cavity; PS, primitive streak; EC, exocoelomic cavity; EPC, ectoplacental cavity

### Whole transcriptome profiling of E5.5 *Mettl14^-/-^* embryos

To investigate the molecular consequences of *Mettl14* depletion in mouse early embryogenesis, we isolated mRNA from control and *Mettl14^-/-^* embryos at E5.5 and performed RNA sequencing. We chose E5.5 mouse embryos because the mutant embryos at this stage were undistinguishable in morphology from the normal littermates, therefore minimizing molecular changes ascribed to secondary developmental defects in the absence of METTL14. The RNA-seq data were mapped to the mouse genome (mm9) with Hisat2, which was published in Nature Protocol to mapping data efficiency. A total of 37979006 (91.4%, WT1), 43551393 (92.32%, WT2) and 46031008 (92.18%, KO1), 62093886 (94.95%, KO2) mapped reads were obtained and used for downstream bioinformatics analysis for the control and *Mettl14^-/-^* embryos, respectively. The results showed that with a cutoff of fold change > 2, P<0.01, there were nearly 1060 differentially expressed genes (DEGs) compared with the control group, indicating that the transcriptome signature was significantly disturbed in E5.5 *Mettl14^-/-^* embryos. Then, replicate Multivariate Analysis of Transcript Splicing (rMATS) software was adopted to analyze the differential alternative splicing events between control and *Mettl14^-/-^* transcriptomes (Fig.3A). We showed that multiple alternative splicing events were affected in the absence of METTL14, with exon skipping to be the most prevalent one (Fig. 3B), which is consistent with previous reports that m^6^A could regulate biological processes through regulating alternative splicing events (Bartosovic et al., 2017). To identify the affected biological processes that may underlie the developmental failure of *Mettl14^-/-^* embryo, gene ontology (GO) analysis was performed with the DEGs. The results showed that dysregulated genes were enriched in embryo development pathways such as in utero embryonic development, anterior/posterior pattern specification, trophectodermal cell differentiation, endoderm development, Wnt signaling pathway, Tgfβ receptor signaling pathway and Notch signaling pathway, all of which are differentiation-related events or signaling pathways essential for mouse early development (Fig. 3C). Thus, our data showed that the early embryonic lethality of *Mettl14* mutants might be a result of impaired differentiation after implantation. Genes, reported function in cell differentiation and anterior/posterior pattern specification, were mapped into an interaction network to illustrate regulatory relationships by Cytocape (Fig. 3D). The genes were significantly dysregulated in *Mettl14-/-* embryos at E5.5. Thus, our study showed that Mettl14-mediated m6A may play a significant role in early embryo development in mice by regulating the expression and alternative splicing of mRNA.

**Figure 3.**
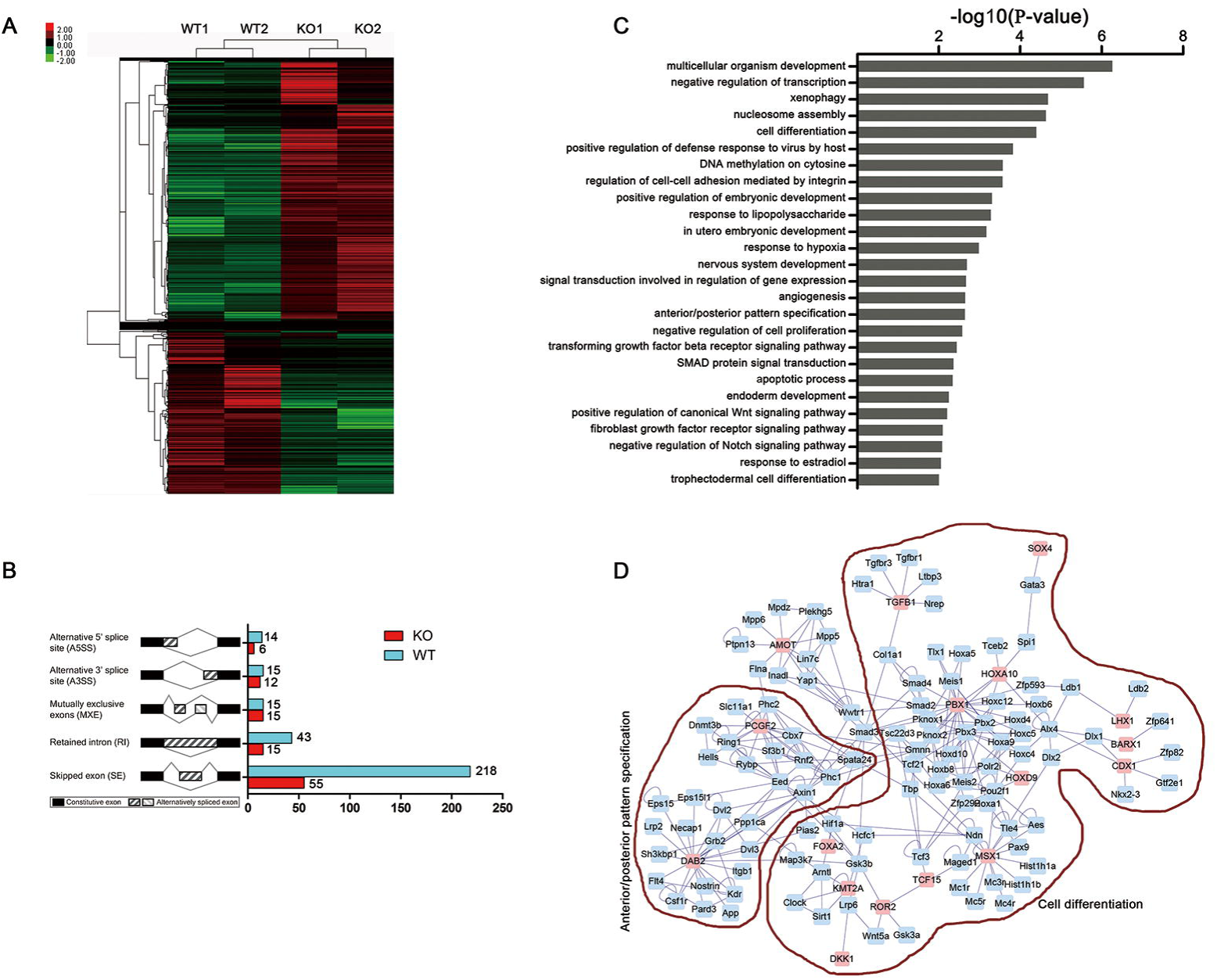
Whole transcriptome analysis of the molecular consequences of *Mettl14* depletion at E5.5. (A) The heat map of differentially expressed genes (Fold change >2, □<0.01) between E5.5 control and *Mettl14^-/-^* embryos. (B) Alternative splicing signature in E5.5 *Mettl14^-/-^* embryos. The rMATS software showed that alternative splicing patterns were disturbed after *Mettl14* depletion, with exon skipping the most profound one. (C) GO analysis of the differentially expressed genes in (A). Development-related events and signaling pathways were overrepresented in the GO terms. (D) The interaction network showing genes involved in anterior/posterior pattern specification and cell differentiation. The regulatory relationships were produced by Cytoscape.

### Depletion of *Mettl14* results in the resistance to differentiation of epiblast *in vivo*

GO analysis indicated that the compromised post-implantation development of *Mettl14^-/-^* embryos might lie in the defects of epiblast differentiation. The conversion of the pluripotent epiblast from a naïve to primed state is an important event essential for mammalian development after implantation. To investigate whether this conversion proceeds normally in the absence of METTL14, we examined the expression of naïve and primed markers in E5.5 normal and *Mettl14^-/-^* embryos with the RNA-seq data. Notably, the expression level of many naïve markers, including *Nr5a2*, *Klf2*, *Rex1* and *Tfcp2l1* in E5.5 *Mettl14^-/-^* embryos were higher compared to the control, while the primed markers, such as *Dnmt3b*, *Otx2*, and *Sox3* were largely down-regulated (Fig. 4A). These data suggested that depletion of *Mettl14* led to impaired epiblast maturation from the naïve to primed state *in vivo*. This result was further validated using immunofluorescent staining of the naïve marker NANOG in E6.5 embryos. While NANOG is expressed in a restricted region in the proximal posterior epiblast in normal E6.5 mouse embryos, its expression was expanded to the whole epiblast after *Mettl14* deletion (Fig. 4C). The expression of general pluripotency marker POU5F1 was little affected (Fig. 4D).

**Figure 4.**
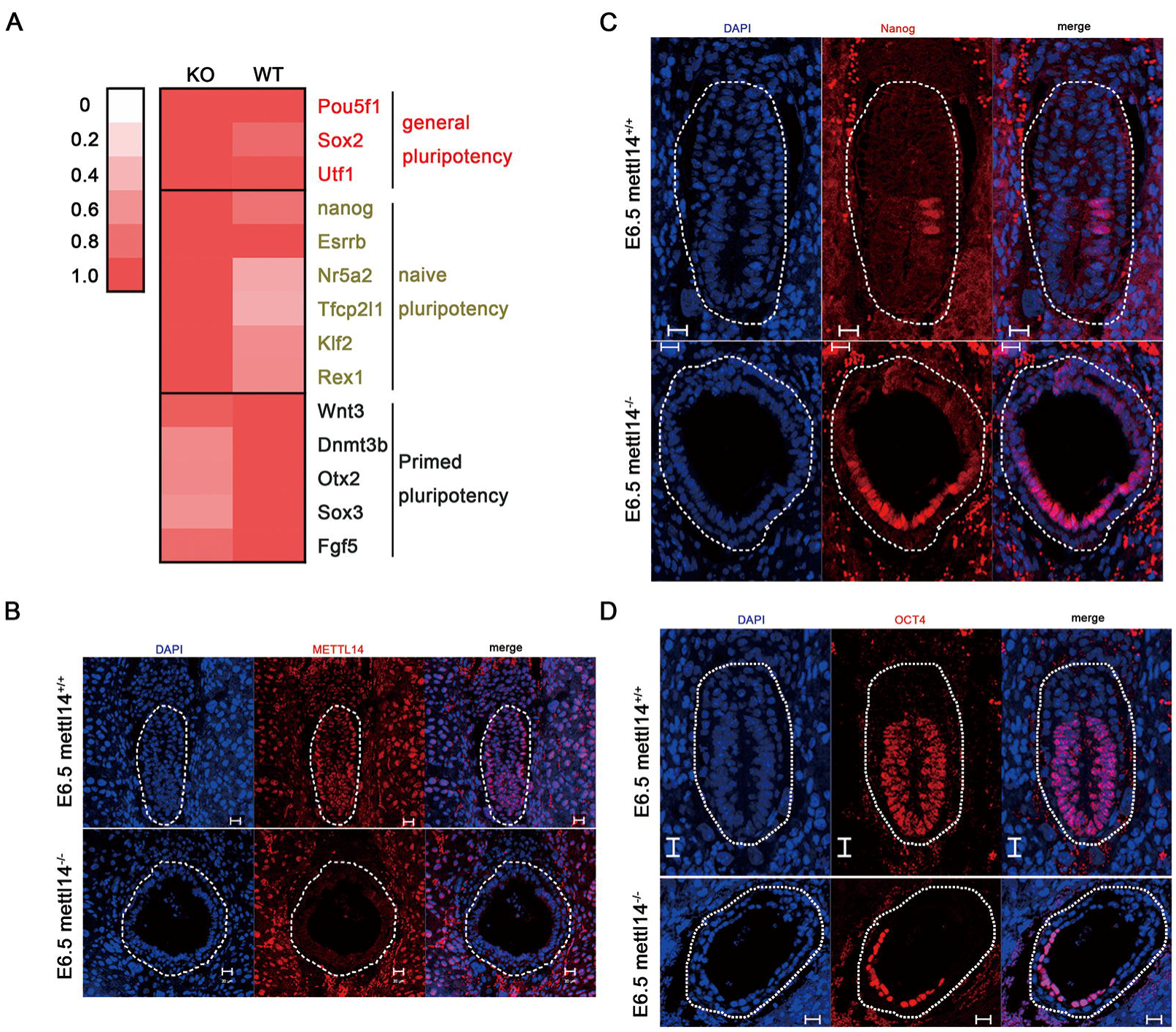
Disruption of *Mettl14* impairs epiblast differentiation *in vivo*. (A) The relative expression of naïve and primed markers in E5.5 control and *Mettl14^-/-^* embryos measured by RNA-seq. The expression level was represented as the ratio of each gene’s FPKM value in control to that in *Mettl14^-/-^* embryos (general and naïve pluripotency markers), or the ratio of each gene’s FPKM value in *Mettl14^-/-^* to that in control embryos (primed markers). (B, C and D) Immunofluorescent staining of METTL14, NANOG and OCT4 in E6.5 control and *Mettl14^-/-^* embryos. Note the failure of NANOG downregulation in the absence of METTL14.

### Mettl14 facilitates the conversion from naïve to primed pluripotency

To better resolve the function of *Mettl14* in mouse early embryogenesis, we attempted to derive ESCs from E3.5 blastocysts of *Mettl14^+/-^* intercrosses for further analysis. *Mettl14^-/-^* blastocysts were largely normal and could not be morphologically distinguished from the normal counterparts. A total of 35 blastocysts from 4 mice were obtained for ESCs derivation, among which 32 colonies were successfully grown out and 8 of them were *Mettl14^-/-^*, as determined with genotyping (Fig.5A, S1). Disruption of *Mettl14* was further confirmed by immunofluorescence and Western blot analysis (Fig. 5B, 5C). While WT ESCs exhibited typical dome-shaped morphology, we observed relatively flattened and irregular morphology of *Mettl14^-/-^* ESCs in 2i/L medium (Fig. 5D). *Mettl14* depletion also decreased ESCs proliferation compared to the control (Fig. 5E). However, the key pluripotency regulators like *Pou5f1*, *Nanog* in *Mettl14^-/-^* ESCs showed comparable protein levels to those in the WT ESCs (Fig. 5F, 5G). Additionally, the activity of alkaline phosphatase (AP), a typical ESCs marker, remained constant in *Mettl14^-/-^* ESCs (Fig.5H). These data suggest that depletion of *Mettl14* has little effect on ESCs self-renewal.

**Figure 5.**
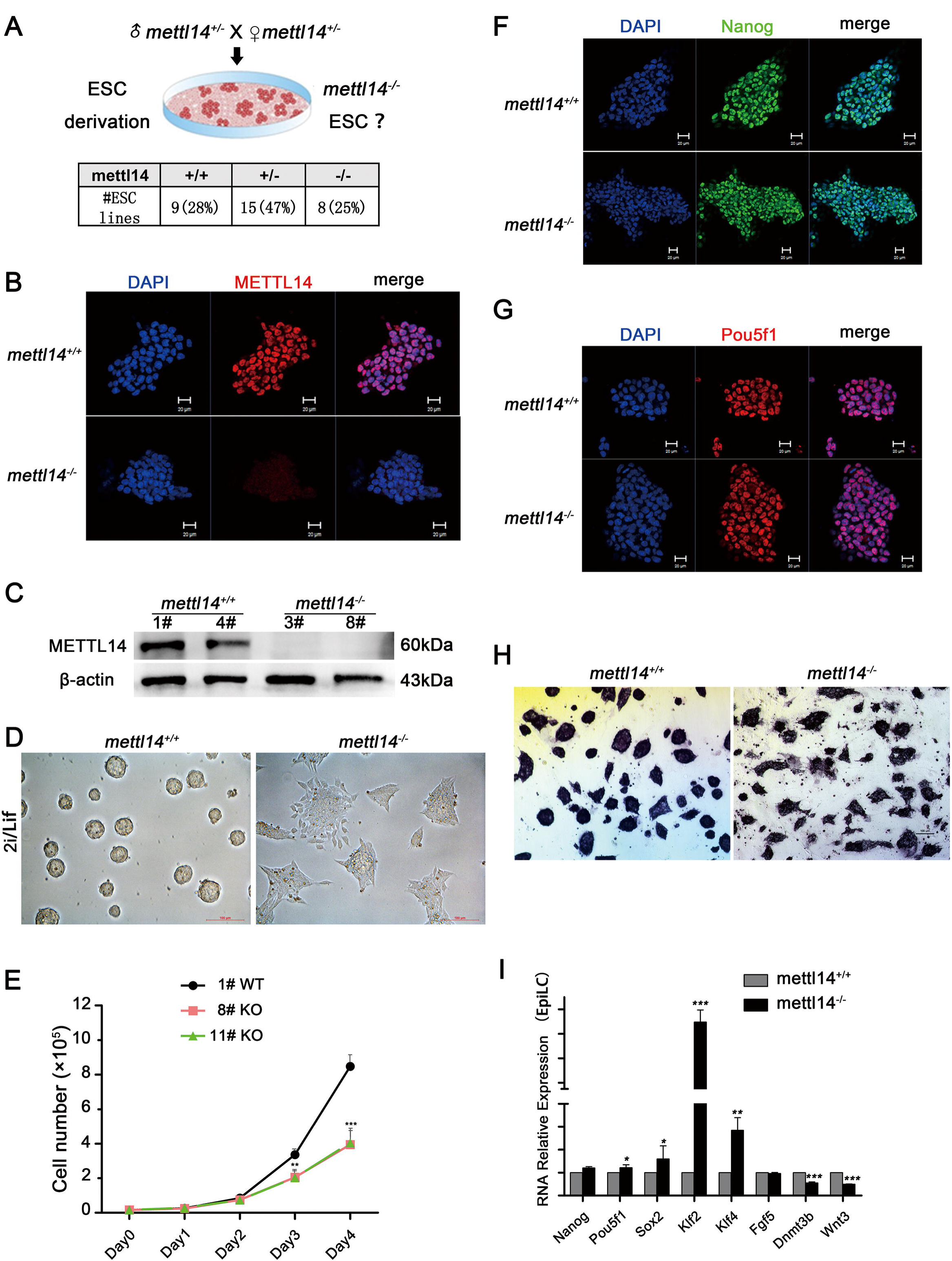
Depletion pf *Mettl14* leads to resistance to conversion from naïve to primed state of mouse embryonic stem cells (ESCs). (A) Scheme showing the derivation of *Mettl14^-/-^* ESCs. *Mettl14^-/-^* ESCs at Mendelian ratio. (B and C) Immunofluorescent staining (B) and Western blot (C) analysis of the expression of METTL14 in control and *Mettl14^-/-^* ESCs. (D) The morphology of control and *Mettl14^-/-^* ESCs. Note the loss of domed shape of *Mettl14^-/-^* ESCs. (E) Proliferation of WT and *Mettl14^-/-^* ESCs. The cumulative population doubling was shown. (F and G) Immunofluorescent staining analysis of the expression of POU5F1 and NANOG in control and *Mettl14^-/-^* ESCs. (H) Alkaline phosphatase (AP) staining of control and *Mettl14^-/-^* ESCs. (I) qRT-PCR analysis of the expression of naïve and primed markers in epiblast like cells.

We further tested the ability of *Mettl14^-/-^* ESCs to convert from naïve pluripotency to the primed state *in vitro*. WT and *Mettl14^-/-^* ESCs were cultured in primed epiblast like cell (EpiLC) medium containing Fgf2 and Activin A for three days (Brons et al., 2007; Hayashi et al., 2011). Both WT and *Mettl14^-/-^* ESCs showed a flattened morphology during the pluripotency conversion and down-regulated naïve markers and up-regulated primed markers. However, the naïve pluripotency markers (*Klf2*, *Klf4*) and the primed pluripotency markers (*Dnmt3b, Wnt3*) were resistant to down-regulation and up-regulation, respectively, compared to the control (Fig. 5I), indicating impaired pluripotency conversion *in vitro*. It is noticeable that the expression of the naïve marker *Nanog* and the primed marker *Fgf5* were comparable to that in the control, suggesting that the conversion from naïve to primed state was initiated but compromised in the absence of METTL14. Thus, loss of *Mettl14* and the resulting decreased m^6^A level hampers the priming and further differentiation competence of ESCs, which might account for the impaired cell fate commitment of the epiblast in vivo.

## DISCUSSION

Eukaryotic mRNA posttranscriptional modification is essential for mRNA maturation, including addition of a 5’ cap, addition of a 3’ poly-adenylation tail, and splicing. Recently, attention has been focused on the physiological function of m^6^A (2-5). m^6^A, the most prevalent internal (non-cap) modification present in mRNA, that affects corresponding mRNA stability and translation status, and is involved in multiple biological processes, including regulation of stem cell differentiation and reprogramming (Batista et al., 2014; Geula et al., 2015; Liu et al., 2014), circadian periods (Fustin et al., 2013), cell cycle, splicing, and embryonic development (Horiuchi et al., 2013; Horiuchi et al., 2006; Ping et al., 2014). However, the function of m^6^A in mammalian early embryo development remains largely unclear. In this study, we found that *Mettl14*, and maybe *Mettl14* mediated m^6^A modification was indispensable for mouse post-implantation embryogenesis by regulating the expression of important regulators for conversion of the epiblast from naïve to the primed state.

Related studies about m^6^A during post-implantation embryo development mainly relied on embryonic stem cells and embryoid bodies, which, after all, are models in vitro. In a previous study, Shay et al reported that *Mettl3* mutant led to embryonic lethality at E6.5 because of failure to down-regulation of *nanog* mRNA. Confusingly, Wang et al showed that m^6^A methylation inversely correlated with mRNA stability and gene expression enriched in developmental regulators. METTTL14, as the core component of the RNA methyltransferase complex, is active and possesses different sets of transcripts compared with METTL3 even though many of their targets overlap. One potential mechanism seems to be the selective regulation of different functions and pathways (Yanan Yue, 2015). Recently, studies have reported that catalysis of the N6-methyl-adenosine mRNA modification m^6^A by a complex containing METTL3 and METTL14 is indispensable for normal spermatogenesis in mice, which showed that Vasa-Cre-mediated ablation of Mettl3 or Mettl14 led to a loss of m^6^A and consequent depletion of spermatogonial stem cells (Lin et al., 2017; Xu et al., 2017). However, Str8-Cre-mediated deletion of either *Mettl3* or *Mettl14* in advanced germ cells did not affect normal spermatogenesis, which indicates independence of METTL3 and METTL14 (Lin et al., 2017).

In our study, we generated *Mettl14* mutant embryos and found that ablation of *Mettl14* led to abnormal embryo development since E6.5. To further investigate the mechanism underlying this phenotype, RNA sequencing (RNA-Seq) based on paired-end high-throughput sequencing methods was adopted to provide insight into the transcriptome of E5.5. Transcriptome analysis revealed that deletion of *Mettl14* resulted in dysregulation of plenty of gene expression enriched in embryo development pathways such as trophectodermal cell differentiation, endoderm development, Wnt signaling pathway, in utero embryonic development, anterior/posterior pattern specification, transforming growth factor beta receptor signaling pathway and Notch signaling pathway, all of which are differentiation related events or signaling pathways essential for mouse early development. However, the previously reported key pluripotency factors such as Nanog do not make a significant difference.

An important question is raised: what is the mechanism responsible for the phenotype caused by *Mettl14* mutation? Whether it is a result of failure to decrease the *Nanog* expression as previously reported in *Mettl3* mutan(Geula et al., 2015) or a result of failure of expression of differentiation-related genes as revealed in our study needs further confirmation. Inconsistent with other naïve markers such as *Klf2*, mRNA level of *Nanog* showed no significant difference compared with the WT group in *Mettl14^-/-^* embryos. It is possible that other regulator may contribute to epiblast maturation, which needs further investigation. The functional mechanisms of m^6^A and its related regulators in post-transcriptional regulation of epiblast maturation is only starting to be uncovered.

## MATERIAL AND METHODS

### Mice

All mouse lines were kept in compliance with the guidelines of the Animal Care and Use Committee of the Institute of Zoology at the Chinese Academy of Sciences. Mice were killed under standard protocols, and all efforts were made to minimize suffering. All mouse strains were maintained in a C57BL/6 background. *Mettl14^+/-^* female mice were mated with *Mettl14^+/-^* male mice to generate *Mettl14^-/-^* embryos. Gestational age of embryos was determined by checking vaginal plugs, with noon of the day of the plug appearance defined as embryonic day (E) E0.5.

### Immunohistofluorescent staining and imaging

For immunohistofluorescent staining, the 5□μm sections of post-implantation embryos were incubated with 5% donkey serum in 0.3% triton X-100 for 1□hr after rehydration and antigen retrieval. Then the sections were incubated with primary antibody overnight at 4°C and then washed in PBS for three times. The membranes were incubated with corresponding F499- or F488-conjugated goat anti-rabbit IgG (1:400, Invitrogen) for 1hr at room temperature. Finally, images were collected using confocal laser scanning microscopy (Carl Zeiss Inc).

### ESCs derivation, culture and conversion of ESCs to EpiLCs

ESCs derivation was performed as previously described with only little modification(Bryja et al., 2006). Briefly, E3.5 blastocysts were obtained and seeded separately on MEF feeders in 3.5 cm culture dishes in KSR ES medium. One week later, the outgrowth of each blastocyst was picked and disaggregated with TypLE Express Enzyme (Gibco), and transferred to 96-well plate in 2i/L medium (N2B27 medium supplemented with 1 μm PD0325901, 3 μm CHIR99021 and 1000 units ml^-1^ LIF). About three days later, clones were disaggregated and transferred to 24-well plates for routine culture. ESCs were routinely maintained on 0.2% gelatin coated dishes in 2i/L medium and propagated at a split ratio of 1:5.

For conversion of ESCs to EpiLCs, 1×10^5^ ESCs were seeded in one well of a 12-well plate coated with Matrigel in N2B27 medium containing Activin A (20 ng/ml), bFGF (12 ng/ml), and 1% KSR for three days. Medium was changed every day.

### mRNA extraction and quantitative real time-PCR

Total RNA was extracted from testis using TRIzol kit following the manufacturer’s instructions. mRNA level of each gene was validated by quantitative real-time PCR (qRT-PCR) analysis (Roche480) according to the manufacturer’s instruction. Primer sets used were listed in Table S1.

### Western blot analysis

A total of 2X10^6^ ESCs per sample were mixed with 2X SDS sample buffer and boiled for 5 minutes at 100°C for SDS-PAGE. Western blotting was performed as described previously, using the primary antibody dilution anti-METTL14 (ATLAS ANTIBODIES, HPA038002) at 1:1000, anti-Nanog (Abcam, ab62734), anti-Pou5f1 (Santa Cruz, sc-5279), anti-β-actin (Zhongshan Golden Bridge Biotechnology) at 1:1000. Secondary antibodies were horseradish peroxidase-linked. Horse-radish peroxidase-linked secondary antibodies (Zhongshan Golden Bridge Biotechnology) were diluted at 1:2000. Protein bands were detected using Thermo Supersignal West Pico chemiluminescent substrate.

### Whole transcriptome profiling

Four E5.5 embryos were subjected to RNA-seq. RNA was isolated from single E5.5 embryo and amplified using SMART-Seq™ v4 Ultra™ Low Input RNA Kit (Clontech), and sequenced by Novogene Corporation. Then we identified the genotype of four embryos based on *Mettl14* qRT-PCR of parts of the returned amplified cDNA. Fortunately, two embryos were identified as *Mettl14^-/-^*, while the other two embryos expressing *Mettl14* mRNA were classified as control group. Library construction was performed following Illumina manufacturer suggestions. Libraries were sequenced on the Illumina Hiseq PE150.

### Statistical analysis

The RNA-seq data were mapped to the mouse genome (mm9) with the software published, and differential expression genes were revealed by using the R package DEseq2. The mapping outcomes were sorted with the software Samtools to obtain a regular processed results; after getting the reads matrix of each group by using Stringtie, differentially expressed genes were screened with the R package (DEseq2). All data presented were collected from at least three independent experiments and analyzed using SPSS (SPSS China). Data were expressed as mean ± SEM and significance of differences was evaluated with Student’s t-test.

## ACKNOWLEDGEMENT

This study was supported by the National Key R&D Program of China (No 2016YFA0100400; 2016YFC1000600) and the National Natural Science Foundation of China (31472055, 31671559), and Youth Innovation Promotion Association CAS (2017114).

---

Figure S1. Genotyping result of the ES colonies.

Table S1. Primer sets used were listed.

## REFERENCES

Arnold, S. J. and Robertson, E. J. (2009). Making a commitment: cell lineage allocation and axis patterning in the early mouse embryo. Nat Rev Mol Cell Biol 10, 91–103.

Bartosovic, M., Molares, H. C., Gregorova, P., Hrossova, D., Kudla, G. and Vanacova, S. (2017). N6-methyladenosine demethylase FTO targets pre-mRNAs and regulates alternative splicing and 3’-end processing. Nucleic Acids Res 45, 11356–11370.

Batista, P. J., Molinie, B., Wang, J., Qu, K., Zhang, J., Li, L., Bouley, D. M., Lujan, E., Haddad, B., Daneshvar, K., et al. (2014). m(6)A RNA modification controls cell fate transition in mammalian embryonic stem cells. Cell stem cell 15, 707–719.

Brons, I. G., Smithers, L. E., Trotter, M. W., Rugg-Gunn, P., Sun, B., Chuva de Sousa Lopes, S. M., Howlett, S. K., Clarkson, A., Ahrlund-Richter, L., Pedersen, R. A., et al. (2007). Derivation of pluripotent epiblast stem cells from mammalian embryos. Nature 448, 191–195.

Bryja, V., Bonilla, S. and Arenas, E. (2006). Derivation of mouse embryonic stem cells. Nat Protoc 1, 2082–2087.

Cong, L., Ran, F. A., Cox, D., Lin, S., Barretto, R., Habib, N., Hsu, P. D., Wu, X., Jiang, W., Marraffini, L. A., et al. (2013). Multiplex genome engineering using CRISPR/Cas systems. Science (New York, N.Y.) 339, 819–823.

Evans, M. J. and Kaufman, M. H. (1981). Establishment in culture of pluripotential cells from mouse embryos. Nature 292, 154–156.

Fustin, J. M., Doi, M., Yamaguchi, Y., Hida, H., Nishimura, S., Yoshida, M., Isagawa, T., Morioka, M. S., Kakeya, H., Manabe, I., et al. (2013). RNA-methylation-dependent RNA processing controls the speed of the circadian clock. Cell 155, 793–806.

Geula, S., Moshitch-Moshkovitz, S., Dominissini, D., Mansour, A. A., Kol, N., Salmon-Divon, M., Hershkovitz, V., Peer, E., Mor, N., Manor, Y. S., et al. (2015). Stem cells. m6A mRNA methylation facilitates resolution of naive pluripotency toward differentiation. Science (New York, N.Y.) 347, 1002–1006.

Hayashi, K., Ohta, H., Kurimoto, K., Aramaki, S. and Saitou, M. (2011). Reconstitution of the mouse germ cell specification pathway in culture by pluripotent stem cells. Cell 146, 519–532.

Horiuchi, K., Kawamura, T., Iwanari, H., Ohashi, R., Naito, M., Kodama, T. and Hamakubo, T. (2013). Identification of Wilms’ tumor 1-associating protein complex and its role in alternative splicing and the cell cycle. J Biol Chem 288, 33292–33302.

Horiuchi, K., Umetani, M., Minami, T., Okayama, H., Takada, S., Yamamoto, M., Aburatani, H., Reid, P. C., Housman, D. E., Hamakubo, T., et al. (2006). Wilms’ tumor 1-associating protein regulates G2/M transition through stabilization of cyclin A2 mRNA. Proceedings of the National Academy of Sciences of the United States of America 103, 17278–17283.

Huang, X., Balmer, S., Yang, F., Fidalgo, M., Li, D., Guallar, D., Hadjantonakis, A. K. and Wang, J. (2017). Zfp281 is essential for mouse epiblast maturation through transcriptional and epigenetic control of Nodal signaling. Elife 6.

Jia, G., Fu, Y., Zhao, X., Dai, Q., Zheng, G., Yang, Y., Yi, C., Lindahl, T., Pan, T., Yang, Y. G., et al. (2011). N6-methyladenosine in nuclear RNA is a major substrate of the obesity-associated FTO. Nature chemical biology 7, 885–887.

Kalkan, T., Olova, N., Roode, M., Mulas, C., Lee, H. J., Nett, I., Marks, H., Walker, R., Stunnenberg, H. G., Lilley, K. S., et al. (2017). Tracking the embryonic stem cell transition from ground state pluripotency. Development 144, 1221–1234.

Lin, Z., Hsu, P. J., Xing, X., Fang, J., Lu, Z., Zou, Q., Zhang, K. J., Zhang, X., Zhou, Y., Zhang, T., et al. (2017). Mettl3-/Mettl14-mediated mRNA N(6)-methyladenosine modulates murine spermatogenesis. Cell Res 27, 1216–1230.

Liu, J., Yue, Y., Han, D., Wang, X., Fu, Y., Zhang, L., Jia, G., Yu, M., Lu, Z., Deng, X., et al. (2014). A METTL3-METTL14 complex mediates mammalian nuclear RNA N6-adenosine methylation. Nature chemical biology 10, 93–95.

Martin, G. R. (1981). Isolation of a pluripotent cell line from early mouse embryos cultured in medium conditioned by teratocarcinoma stem cells. Proceedings of the National Academy of Sciences of the United States of America 78, 7634–7638.

Ping, X. L., Sun, B. F., Wang, L., Xiao, W., Yang, X., Wang, W. J., Adhikari, S., Shi, Y., Lv, Y., Chen, Y. S., et al. (2014). Mammalian WTAP is a regulatory subunit of the RNA N6-methyladenosine methyltransferase. Cell Res 24, 177–189.

Sibbritt, T., Patel, H. R. and Preiss, T. (2013). Mapping and significance of the mRNA methylome. Wiley Interdiscip Rev RNA 4, 397–422.

Tesar, P. J., Chenoweth, J. G., Brook, F. A., Davies, T. J., Evans, E. P., Mack, D. L., Gardner, R. L. and McKay, R. D. (2007). New cell lines from mouse epiblast share defining features with human embryonic stem cells. Nature 448, 196–199.

Wang, Y., Li, Y., Toth, J. I., Petroski, M. D., Zhang, Z. and Zhao, J. C. (2014). N6-methyladenosine modification destabilizes developmental regulators in embryonic stem cells. Nat Cell Biol 16, 191–198.

Xu, K., Yang, Y., Feng, G. H., Sun, B. F., Chen, J. Q., Li, Y. F., Chen, Y. S., Zhang, X. X., Wang, C. X., Jiang, L. Y., et al. (2017). Mettl3-mediated m(6)A regulates spermatogonial differentiation and meiosis initiation. Cell Res 27, 1100–1114.

Yanan Yue, J. L. a. C. H. (2015). RNA N6-methyladenosine methylation in post-transcriptional gene expression regulation. GENES&DEVELOPMENT.

Zheng, G., Dahl, J. A., Niu, Y., Fedorcsak, P., Huang, C. M., Li, C. J., Vagbo, C. B., Shi, Y., Wang, W. L., Song, S. H., et al. (2013). ALKBH5 is a mammalian RNA demethylase that impacts RNA metabolism and mouse fertility. Molecular cell 49, 18–29.

